# Natural sleep, but not propofol-induced anesthesia preserves spatial and procedural memory consolidation

**DOI:** 10.64898/2026.09.05.749587

**Authors:** Eva-Maria Kurz, Annika Hanert, Axel Fudickar, Maike Dittmar, Fee Hoppe, Julius Rave, Sarah Philippen, Frederik D. Weber, Robby Schönfeld, Svenja Brodt, Jan Born, Thorsten Bartsch

**Author notes:** these authors contributed equally (Eva-Maria Kurz, Annika Hanert). Corresponding author/Lead Author: Thorsten Bartsch, MD Memory Disorders and Plasticity Group Dept. of Neurology University Hospital Schleswig-Holstein, Kiel Arnold-Heller-Str. 3 24105 Kiel, Germany.

## Abstract

Propofol-induced general anesthesia and natural sleep are both associated with GABA-A-ergic inhibition and slow-wave activity. Whereas sleep actively supports memory consolidation, consolidation may be disrupted during anesthesia. Here, we compared EEG and memory data from 19 participants undergoing surgery under propofol-induced anesthesia with data from 17 participants taking a nap. In both groups, participants completed a hippocampus-dependent spatial memory task (Virtual Water Maze) and a procedural memory task (Mirror Tracing) before (pre) and after (post) an interval filled with either anesthesia or sleep. Performance declined from pre to post in the propofol group for both tasks, whereas it remained stable in the sleep group. Slow oscillations (SO) occurred during both non-REM sleep and propofol-induced anesthesia. However, only during non-REM sleep did SOs show increased spindle activity during the SO upstate. Together, these findings suggest that propofol anesthesia disrupts memory consolidation, most likely because SOs under anesthesia lack the spindle coupling that mediates the hippocampal-neocortical information transfer underlying systems memory consolidation.

## Introduction

Sleep and general anesthesia are neurophysiologically distinct states, despite some similarities in their electroencephalographic (EEG) signatures and a shared dependence on GABA-A-ergic inhibition (Akeju & Brown, 2017). While natural sleep is homeostatically regulated with the brain cycling through non-rapid eye movement sleep (N1-N3, non-REM) and REM sleep in an ultradian rhythm (Borbely et al., 2016; Cajochen et al., 2024), general anesthesia is a clinically induced state of unconsciousness. Propofol enhances GABA-A-mediated inhibition and thereby alters neuronal excitability across cortical and subcortical networks; like non-REM sleep, it can induce slow oscillations (SO; approximately 0.5–1.5 Hz) and slow-wave activity, but the underlying circuit dynamics are not identical and the inhibitory pattern appears to be more global than during sleep (Bonhomme et al., 2019; Purdon et al., 2015; Vanini et al., 2020).

A key electrophysiological distinction is that propofol-induced anesthesia typically produces an anterior alpha rhythm in the 8–12 Hz range (Purdon et al., 2015), whereas N2 sleep is characterized by sleep spindles, thalamocortical waxing and waning oscillations in the approximately 12–15 Hz range (Fernandez & Lüthi, 2020). In sleep, these spindle events are not merely an epiphenomenon of reduced arousal but form part of a highly coordinated replay architecture in which hippocampal ripples, sleep spindles, and SOs interact to support the transfer and redistribution of newly encoded information from hippocampal to neocortical networks (Brodt et al., 2023; Clemens et al., 2007; Staresina et al., 2015). The precise nesting of fast sleep spindles into the depolarized SO upstate has been repeatedly associated with overnight memory consolidation in declarative (Bastian et al., 2022; Hahn et al., 2020; Muehlroth et al., 2019) but also in procedural tasks (Mikutta et al., 2019; Solano et al., 2022) suggesting a contribution of the hippocampus also in offline motor memory consolidation (Baena et al., 2024; Schapiro et al., 2019). By contrast, propofol-induced SOs do not appear to recruit spindle activity in the same way as natural sleep. Propofol slow waves can resemble sleep slow waves in their gross morphology and cortical origin, but they fail to effectively entrain spindle activity and show altered large-scale organization (Murphy et al., 2011). This lack of SO-spindle coupling is particularly important because it suggests that the emergence of slow oscillatory activity under propofol does not necessarily imply a sleep-like functional state. Rather, propofol may generate oscillatory patterns that are superficially similar to those of non-REM sleep but do not effectively support the coordinated hippocampal-neocortical communication required for memory stabilization.

At the cellular level, propofol has been shown to impair hippocampal plasticity, including long-term potentiation in CA1, indicating that its effects are not limited to global arousal suppression but extend to synaptic mechanisms underlying to memory formation (Takamatsu et al., 2005; Wei et al., 2002). In humans, sedative levels of propofol suppressed hippocampal responses and diminished memory performance in comparison to a placebo control croup (Pryor et al., 2015). The early period after learning appears to be particularly sensitive to the effect of propofol as the recall of word lists was only impaired when propofol-induced general anesthesia was induced within around 13 minutes after learning whereas there was no effect when anesthesia was induced approximately 105 minutes after learning (Moon et al., 2020). Similarly, Iggena et al. (2022) found impaired spatial navigation after anesthesia only when memories were encoded directly before anesthesia and with encoding more than 60 minutes before anesthesia induction.

Despite these advances, sleep and propofol-induced general anesthesia have not yet been directly compared within the same experimental framework with respect to both memory performance and EEG oscillatory dynamics. This gap is particularly relevant because the mechanistic interpretation of propofol-induced SOs remains unresolved. If these oscillations reflect a sleep-like consolidation state, one would expect them to support spindle coupling and memory stabilization. If, however, they represent a dysfunctional state of oscillatory activity, then one would predict preserved or even enhanced SO generation without the coordinated spindle recruitment necessary for consolidation. In the present study, we compared memory performance and EEG activity during propofol-induced anesthesia and natural sleep in two hippocampus-associated tasks: a spatial memory task and a procedural memory task. We hypothesized that sleep, but not propofol-induced anesthesia, would support consolidation across the retention interval. Furthermore, we predicted that SOs during sleep would be accompanied by spindle activity during the SO upstate, whereas SOs during propofol anesthesia would lack this coupling and would therefore be dissociated from successful memory consolidation (Murphy et al., 2011).

## Methods

### Participants

A total of 39 adults participated in the study. Of these, 19 patients of the Ear, Nose and Throat (ENT) Department, University Hospital Schleswig-Holstein, Campus Kiel scheduled for ENT surgery under general anesthesia using propofol were recruited as the propofol group and 20 participants were recruited as a sleep control group. Before the start of the actual study, a pilot study was conducted including four participants to test the feasibility of continuous EEG recording under anesthesia during ENT surgery. Once the results were satisfactory, the actual study began. The study was approved by the local ethics committee, and all participants gave informed consent before inclusion. All procedures were in accordance with the Declaration of Helsinki.

Of the 19 participants in the propofol group (mean age 29.32 ± 2.04 years, range 20 – 46, 11 female), two participants were not included in the analysis of EEG data due to poor signal quality, and three to six participants (depending on the task) did not provide data in the memory task. The sleep group consisted of 20 university students. Due to insufficient sleep during the nap opportunity (less than 20min non-REM sleep), three participants had to be excluded from the sleep group, leaving 17 participants (mean age 23.77 ± 0.42 years, range 22 – 29 years, 9 female) for further analyses. Participant numbers included in each analysis are indicated in the tables depicting means and standard errors (Table 1 and Table S1-S3).

**Table 1.** Characteristics of sleep and anesthesia (M ± SEM)

|  | Sleep<br>(n = 17) | Propofol<br>(n = 17) |
| --- | --- | --- |
| TST/LOC (min) | 90.09 ± 3.87 | 107.26 ± 11.04 |
| N1 (min) | 8.56 ± 1.20 | - |
| N2 (min) | 44.56 ± 3.52 | - |
| N3/SWS (min) | 27.00 ± 3.55 | - |
| REM (min) | 9.97 ± 2.58 | - |
| WASO (min) | 8.24 ± 2.15 | - |
| SO characteristics |  |  |
| density (event/min) |  |  |
| frontal* | 3.77 ± 0.22 | 3.98 ± 0.19 |
| central | 3.22 ± 0.23 | 4.03 ± 0.19 |
| amplitude (μV) | | |
| frontal* | 205 ± 7.94 | 153 ± 11.11 |
| central | 152 ± 5.80 | 155 ± 9.32 |
| slope ($\mu\text{V/s}$) | | |
| frontal* | $649 \pm 34.46$ | $316 \pm 31.40$ |
| central | $413 \pm 19.68$ | $303 \pm 22.96$ |
| Dominant Frequency |  |  |
| frontal* | $12.75 \pm 0.32$ | $11.25 \pm 0.18$ |
| central | $13.58 \pm 0.27$ | $11.24 \pm 0.18$ |

TST = total sleep time; LOC = loss of consciousness; REM = Rapid Eye Movement; WASO = wake after sleep onset.
\* only 16 participants from the propofol group

Neuropsychological testing did not indicate substantial differences between the sleep and propofol group (see Table S1). Testing included the Rey-Auditory Verbal Learning Test (RAVLT) to assess verbal memory (Rey, 1941), as well as the Trail Making Test A (TMT A) and B (TMT B) to measure attention and cognitive flexibility (Reitan, 1979). Working memory was tested using the digit span task as part of the Wechsler Adult Intelligence Scale (Wechsler, 1997), followed by a word fluency task (Aschenbrenner et al., 2000). There were no group differences in the RAVLT, Trail Making Test and the digit span task. Further, there were no group differences in lexical word fluency, but participants from the propofol group named less words from categories compared to the sleep group (*p* = .007; see also Table S1).

Participants undergoing anesthesia had to fulfil the following inclusion criteria: (i) Elective surgery in the ENT Department, University Hospital Schleswig-Holstein, Campus Kiel under general anesthesia, (ii) ASA status I or II according to the classification of the American Society of Anesthesiologists (ASA) into one of four stages based on their physical condition before anesthesia (I = healthy or II = mild systemic disease); and (iii) body mass index between 20 and 28 kg/m^2^. Exclusion criteria were diagnoses of chronic pain or pre-existing neurological diseases as well as arterial hypertension.

Participants in the sleep group were instructed to refrain from alcohol on the day before and the day of the experiment, to avoid coffee and additional naps on the experimental day, and to sleep only 6 h and rise early on the day of testing.

### Procedure

All participants completed two test sessions, one before and one after the retention interval, during which they performed a virtual water maze (VWM) task and a visuo-motor mirror tracing task (MTT). In the sleep group, the first test session (pre) started around 12:00 pm, and was followed by a 90-minute nap opportunity accompanied by polysomnography in the sleep laboratory of the Department of Neurology of the University of Kiel. The propofol group performed the first test session in the morning of the day of the surgery with variable times between the first test session and start of anesthesia.

The second test session (post) was completed after sleep (3.10 ± 0.06 hours after encoding, range: 2.8 to 3.65 hours) or anesthesia (25.07 ± 5.13 hours after encoding, range: 8 to 70 hours after learning). The propofol group completed the post session at a later time point than the sleep group, due to the necessary recovery period following surgery and anesthesia, which delayed readiness to perform the experimental tasks.

Before the experimental day, participants in the sleep group completed an adaptation nap and neuropsychological assessment. In the propofol group, the neuropsychological assessment was performed the day before surgery. Details on induction and maintenance of anesthesia are reported in the supplementary material.

### Electroencephalographic recording and analyses

EEG recordings during sleep and anesthesia were performed using a portable device, SomnoScreen EEG 10-20 (Somnomedics, Randersacker, Germany). Analyses of non-REM sleep and propofol-induced anesthesia comprised the calculation of power spectra and detection of SOs. Cross-frequency coupling between slow oscillatory activity and activity in the spindle frequency range was evaluated by calculating time frequency representations (TFR) of the SO trough-locked data and calculating the Synchronization Index (Cohen, 2008). Please refer to the supplementary material for further information.

### Virtual water maze

The VWM is based on the hidden platform paradigm in the Morris water maze, a standard test of spatial learning and memory in rodents (Morris, 1984) and humans (Hanert et al., 2024; Schoenfeld et al., 2014). Similar to the hidden platform in the Morris water maze, a hidden treasure box has to be found (Figure 1). During the task, the location of the target is not visible from the first-person view until the participant navigates to a place that is very close to the hidden target. The circular island is sectioned into four quadrants that are marked by different distal cues (i.e. four landmarks: lighthouse, windmill, sailing yacht, and sea buoy). In this study, the target was located in the area of the lighthouse. Participants navigated across the island using a joystick. The first-person view within the virtual reality provided natural conditions to make the outcome comparable with real life conditions. The paradigm presupposes the ability of navigating through a natural environment after its acquisition and tests the performance of place learning.

**Figure 1.**
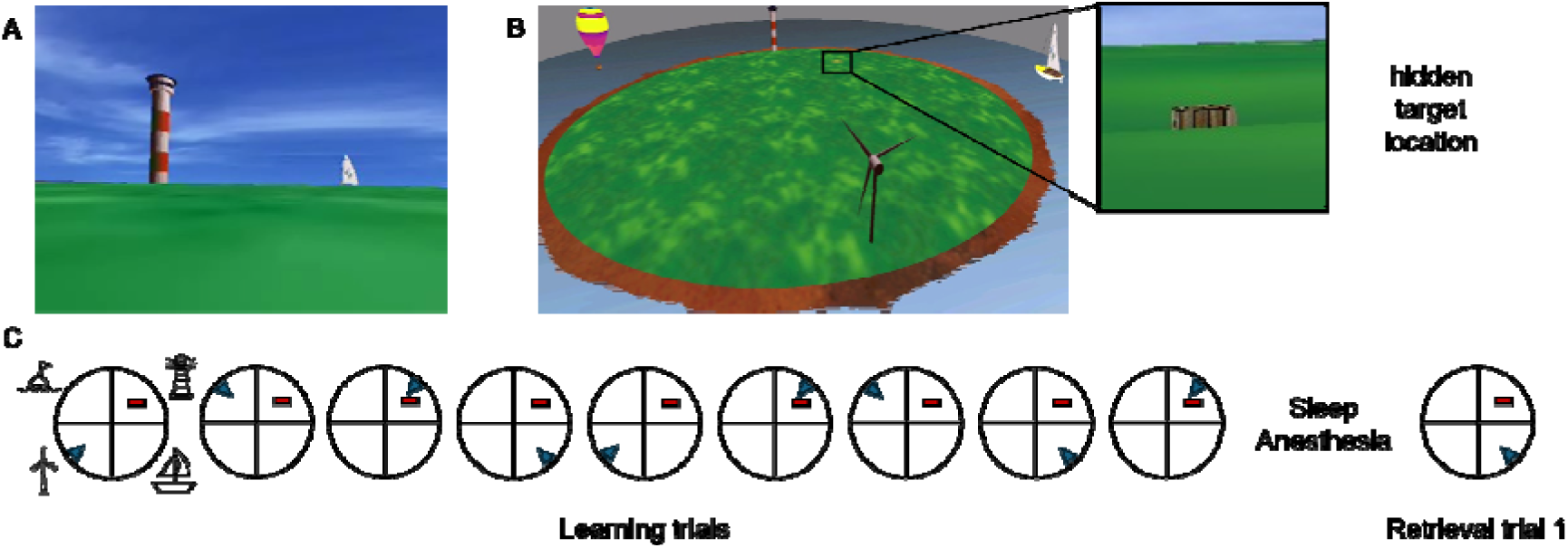
Overview of the Virtual water maze (VWM) task. **A:** Illustration of the island from the participant’s perspective. **B**: Illustration of the island in bird’s-eye view. Participants navigated a circular island from a first-person perspective using a joystick. Four distal landmarks, including a lighthouse, windmill, sailing yacht, and sea buoy, provided stable spatial cues. The hidden treasure chest was located in the quadrant associated with the lighthouse and became visible only when participants approached the target location closely. **C:** Schematic representation of the VWM trial structure. The arrow marks the respective starting position of the participant; the rectangle marks the treasure chest. During the pre-session, participants completed nine learning trials. The first trial included an additional cue in the form of a hot-air balloon above the target, whereas the remaining eight trials required navigation to the same hidden location from different starting positions using only distal cues.

During the first test session (pre sleep or anesthesia), participants completed nine learning trials. They were asked to navigate across an island to find the hidden treasure box and have to remember its location. The first of the nine learning trials was additionally cued with a hot-air balloon placed above the target treasure chest. In the remaining eight learning trials the participants navigated to the same target location but from different starting points using only the distal cues.

In the second test session (post), participants completed four retrieval trials. In the first and second retrieval trial, the target was located at the same position as in the learning trials, with all four distal cues provided in the first but only two distal cues provided in the second retrieval trial. In the remaining two retrieval trials, the target was located opposite to its original position. For the current question of interest only the first retrieval trial was considered.

Performance was evaluated by three measures: latency in seconds, i.e. the duration from starting point to finding the target; path length (divided by pool diameter; in pixels), i.e. the distance moved between the starting point and the target. Both shorter latencies and path lengths are an indication of learning. Lastly, relative dwell time in the target quadrant was calculated. The longer participants dwelled near the target, the better the memory of the position.

Consolidation was evaluated by comparing the first retrieval trial to the eighth learning trial, which was the last learning trial having the same starting position as retrieval trial one.

### Mirror tracing task

The mirror tracing task is a standard non-declarative learning task for procedural visuo-motor learning (Gais & Born, 2004; Groch et al., 2013). The participants were asked to trace a star consisting of five points and bordered by an outer line and an inner line (0.5 cm distance) as fast and accurately as possible. The star is only visible through a mirror, and the starting point is marked at the tip of the uppermost point. Touching or passing the border lines and interrupting the drawn line is counted as an error. Participants performed the task two times, once before and once after the sleep/anesthesia interval. The number of errors and time to complete the star were calculated and compared.

### Statistical analysis

Statistical analyses were done in R version 4.4.1, mainly applying linear mixed-effects models using the library *lme4* (Bates et al., 2015). The R library *parameters* (Lüdecke et al., 2020) was used to obtain model parameters with p-values and degrees of freedom estimated via Satterthwaite approximation. Post-hoc simple slope analyses and pair-wise comparisons were done using the *emmeans* package (Lenth, 2023), correcting for multiple comparisons using Bonferroni method. Models investigating the performance during learning trials in the VWM task included the fixed effects group (propofol/sleep) and learning trials (1 to 9, continuous) and random intercepts for participants. To improve model fit, latency and path length were log-transformed. For both tasks, analysis of group differences in performance across the delay included the fixed effect group (propofol/sleep), time (pre/post) and random intercepts for participants. Analyses of SO characteristics and spectral measures included the fixed effects group (propofol/sleep) and site (frontal/central) and random intercepts for participants. Due to a greater age range in the propofol group (though there was no significant difference between the groups: *z* = -1.26, *p* = .214), age was added as a covariate in all models. The duration between encoding and retrieval was added to all models analyzing memory performance across the sleep/anesthesia interval.

Statistical analysis of TFRs was done using the FieldTrip toolbox (Oostenveld et al., 2011). Separately for the sleep and propofol group baseline-normalized TFRs (average of frontal channels) were tested against zero using dependent sample t-tests and corrected for multiple comparisons using cluster-based permutation tests (Maris & Oostenveld, 2007), with 5000 permutations and a cluster alpha-level of 0.05 (two-tailed), Monte Carlo method. In a second step, baseline-normalized TFRs of the sleep and propofol group were compared using independent sample t-tests, similarly using cluster-based permutation for multiple comparison correction.

## Results

### Spatial navigation

Hippocampus – dependent spatial memory performance was assessed in the Virtual Morris Water Maze by analyzing latency (the duration until reaching the target), path length (from start target position) and relative dwell time in the target quadrant. Latency (Figure 2A) improved across learning trials in both groups, although participants in the propofol group showed a greater improvement than those of the sleep group (learning trial × group interaction: *b* = 0.04, SE = 0.01, 95% CI [0.01, 0.06], *t*(263) = 2.48, *p* = .014; simple slopes propofol group: *b* = -0.06, SE = 0.01, *t*(240) = -5.42, *p* < .001, simple slopes sleep group: *b* = -0.02, SE = 0.01, *t*(240) = -2.43, *p* = .031). Similarly, path length shortened across trials (main effect learning trial: *b* = -0.03, SE = 0.008, 95% CI [-0.04, -0.01], *t*(263) = -3.55, *p* < .001), although here the interaction was not significant (learning trial × group interaction: *b* = 0.03, SE = 0.02, 95% CI [<0.01, 0.06], *t*(263) = 1.92, *p* = .056) and there was no main effect of group (*b* = 0.04, SE = 0.11, 95% CI [-0.17, 0.26], *t*(263) = 0.41, *p* = .683). Relative dwell time similarly improved across learning trials (*b* = 0.01, SE = 0.003, 95% CI [0.01, 0.02], *t*(263) = 4, *p* < .001) but did not differ between groups (*b* = -0.10, SE = 0.05, 95% CI [-0.21, 0.01], *t*(263) = -1.80, *p* = .073) and there was no interaction between group and learning trial (*b* = -0.004, SE = 0.01, 95% CI [-0.02, 0.01], *t*(263) = -0.60, *p* = .548). The participant’s age did not predict these performances in any model (all *p* > .238). See also Table S2 for means and standard errors.

**Figure 2.**
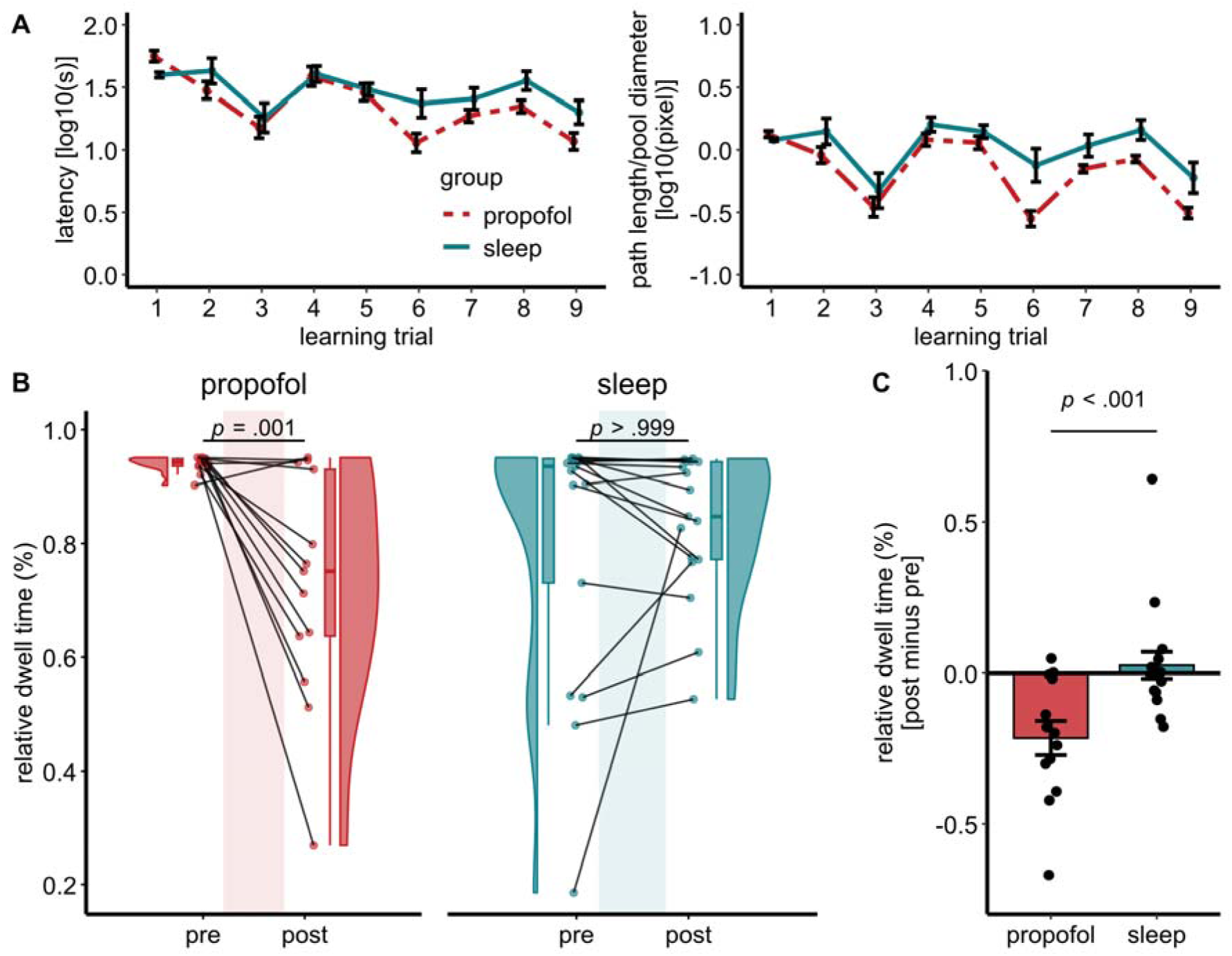
Performance in the virtual water maze task. **A.** Learning performance across the nine acquisition trials, shown as latency (in seconds) to reach the hidden target and path length from start position to target. Both measures improved across trials, indicating acquisition of the task. Data are shown as group means with standard errors. **B.** Spatial memory consolidation across the retention interval, quantified as relative dwell time in the target quadrant during learning trial 8 (pre) and retrieval trial 1 (post). Depicted are density plots, boxplots and individual data points. **C.** Spatial memory consolidation depicted as the difference between retrieval trial 1 (post) and learning trial 8 (pre). Individual data points, groups means and standard errors are shown.

To evaluate spatial memory across the sleep/anesthesia interval, we analyzed the relative dwell time (Figure 2B). Performance deteriorated across the retention interval in the propofol group (*t*(30) = 4.17, *p* = .001), while it stayed constant in the sleep group (*t*(30) = - 0.58, *p* > .999; pre/post × group interaction: *b* = -0.24, SE = 0.07, 95% CI [-0.38, -0.10], *t*(52) = -3.52, *p* < .001). Importantly, this was independent of the amount of time passed between the pre and post test session (*p* = .438) or the participants’ age (*p* = .293). Excluding participants with below-chance performance (< 25% dwell time in the target quadrant) in one of the last three learning trials left the results unchanged (group × time interaction: *b* = -0.20, SE = 0.06, 95% CI [-0.32, -0.09], *t*(48) = -3.53, *p* < .001). The same effects were observed for the latency (group × time interaction: *b* = 0.30, SE = 0.09, 95% CI [0.12, 0.48], *t*(52) = - 3.38, *p* = .001) and path length (group × time interaction: *b* = 0.25, SE = 0.09, 95% CI [0.08, 0.42], *t*(52) = 2.90, *p* = .005).

### Mirror tracing task

Non-declarative procedural memory was assessed using the mirror tracing task. Despite an association of more errors with higher age of the participants (*b* = 11.12, SE = 3.97, 95% CI [3.16, 19.08], *t*(52) = 2.80, *p* = .007), the propofol group did not change in their performance (*t*(30) = 0.88, *p* > .999) while the sleep group made less errors after the retention interval (*t*(30) = 4.99, *p* < .001; pre/post × group interaction: *b* = 15.61, SE = 5.94, 95% CI [3.69, 27.53], *t*(52) = 2.63, *p* = .011, see also Figure 3 and Table S3). There was no effect of the duration of time passed between the immediate and delayed test (*p* = .164). For the duration of task completion there was no interaction between time and group (*p* = .699). The task was generally performed faster after the retention interval (main effect pre/post: *b* = 38.42, SE = 8.01, 95% CI [22.35, 54.49], *t*(52) = 4.80, *p* < .001) and older participants took a longer time (*b* = 18.87, SE = 6.55, 95% CI [5.73, 32.01], *t*(52) = 2.88, *p* = .006).

**Figure 3.**
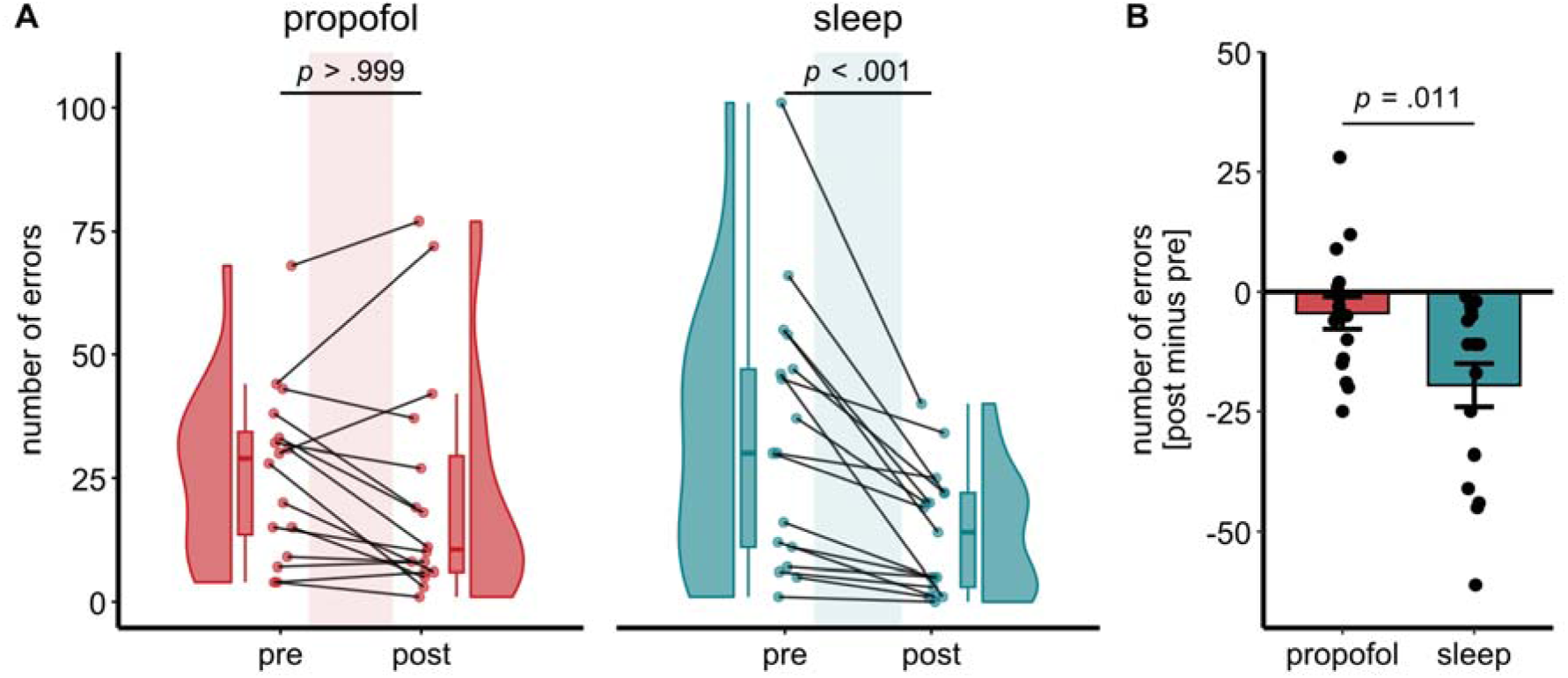
Performance in the procedural visuo-motor learning in the mirror tracing task. **A.** Number of tracing errors before and after the retention interval. Depicted are density plots, boxplots and individual data points. **B.** Procedural memory consolidation depicted as the difference between after (post) and before (pre) the retention interval. Individual data points, groups means and standard errors are shown.

### Sleep and anesthesia

Information on sleep stage durations for the sleep group and duration of loss of consciousness for the propofol group are provided in Table 1. As expected, alpha activity during loss of consciousness was increased in the propofol group and spindle activity during non-REM sleep was increased in the sleep group (see Figure 4).

**Figure 4.**
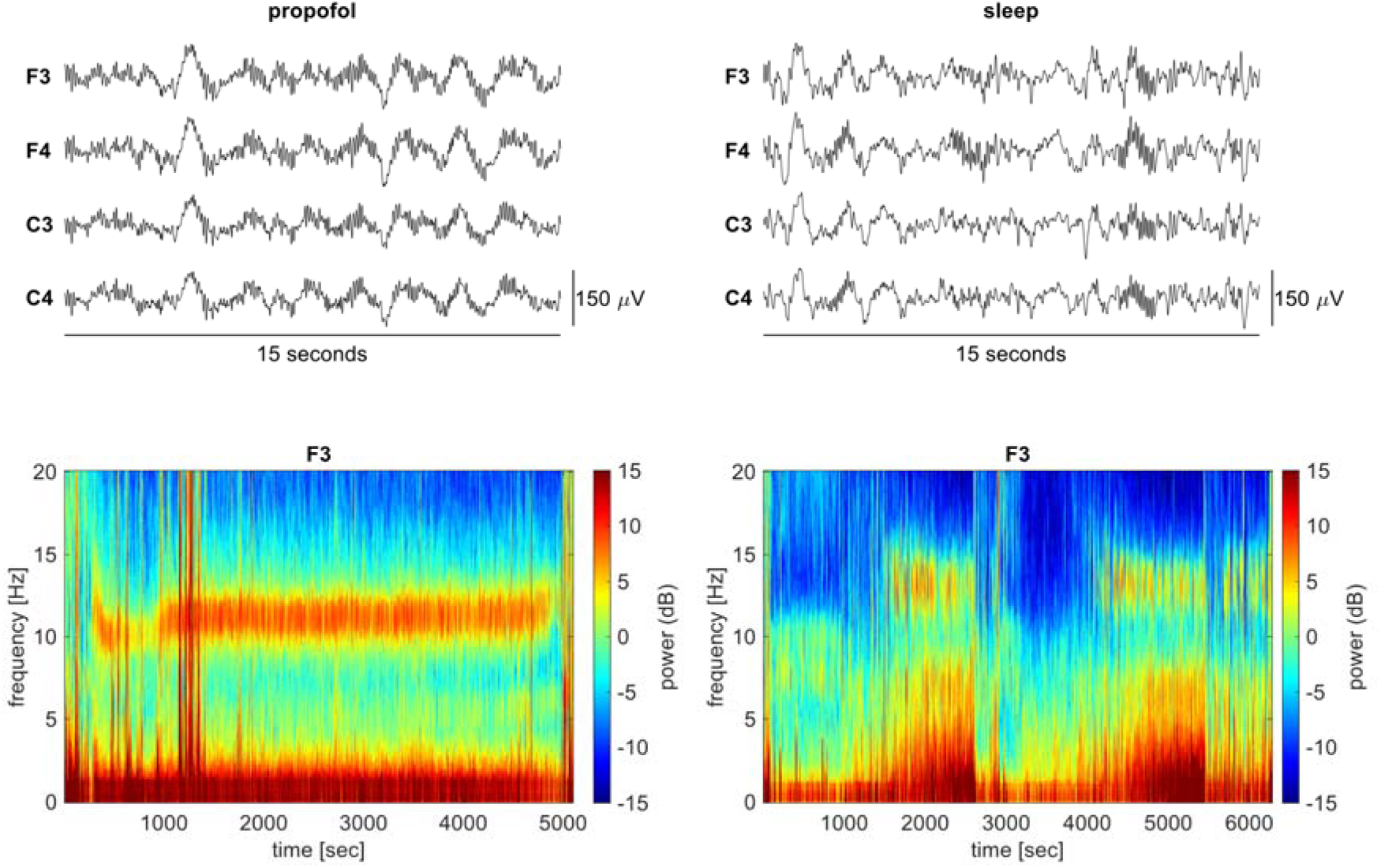
Electrophysiological activity during propofol-induced anesthesia (left column) and non-REM sleep (right column). Top row: Representative 15-s EEG segments illustrating the corresponding state-dependent oscillatory patterns. During propofol anesthesia the signal is dominated by continuous alpha and slow-wave activity, whereas non-REM sleep is characterized by discrete spindles and slow oscillations. Bottom row. Exemplary spectrogram from one participant in the propofol group (left) and one participant in the sleep group (right), for channel F3, similarly depicting continuous alpha activity (∼11 Hz) during propofol-induced anesthesia and discrete spindle events (> 12 Hz) during non-REM sleep.

### Oscillatory activity during sleep and propofol

The dominant (oscillatory) frequency in the power spectra was higher during sleep and had a higher peak power than during propofol at frontal and central sites. Additionally, in the sleep group only, the frequency peak was higher centrally compared to frontally (sleep group: *t*(33.1) = -5.54, *p* < .001; propofol group *p* > .999; group × channel interaction: *b* = 0.81, SE = 0.21, 95% CI [0.38, 1.24], *t*(61) = -3.77, *p* < .001). While in the propofol group the frequency power peak (11 Hz) resembled the characteristic frequency of alpha oscillations, the frequency power peak in the sleep group is characteristic of sleep spindles, that are typically at a lower oscillatory frequency in frontal areas (frontal: 12.75 Hz, central: 13.58 Hz, see Figure 4 and Table 2).

### Slow Oscillations

All three SO characteristics (density, amplitude and slope) differed depending on group and channel (see also Table 1 and Figure 5). SO density, amplitude and slope decreased from frontal to central sites in the sleep group, which was not the case in the propofol group. Regarding site-specific group differences, SO density was greater in the propofol than the sleep group only centrally (*t*(38.4) = 2.88, *p* = .026) but not frontally (*t*(38.9) = 0.86, *p* > .99; group × channel interaction: *b* = -0.57, SE = 0.14, 95% CI [-0.86, -0.28], *t*(61) = -3.98, *p* < .001). Furthermore, SO amplitudes were higher in the sleep than the propofol group frontally (*p* < .001) but not centrally (*p* > .999; group × channel interaction: *b* = -51.67, SE = 6.21, 95% CI [-64.09, -39.26], *t*(61) = -8.33, *p* < .001). Lastly, the SO slope was steeper in the sleep than the propofol group at both sites, but the difference between the groups was greater frontally than centrally (group × channel interaction: *b* = -216.4, SE = 23.86, 95% CI [-264, - 169], *t*(61) = -9.07, *p* < .001).

**Figure 5.**
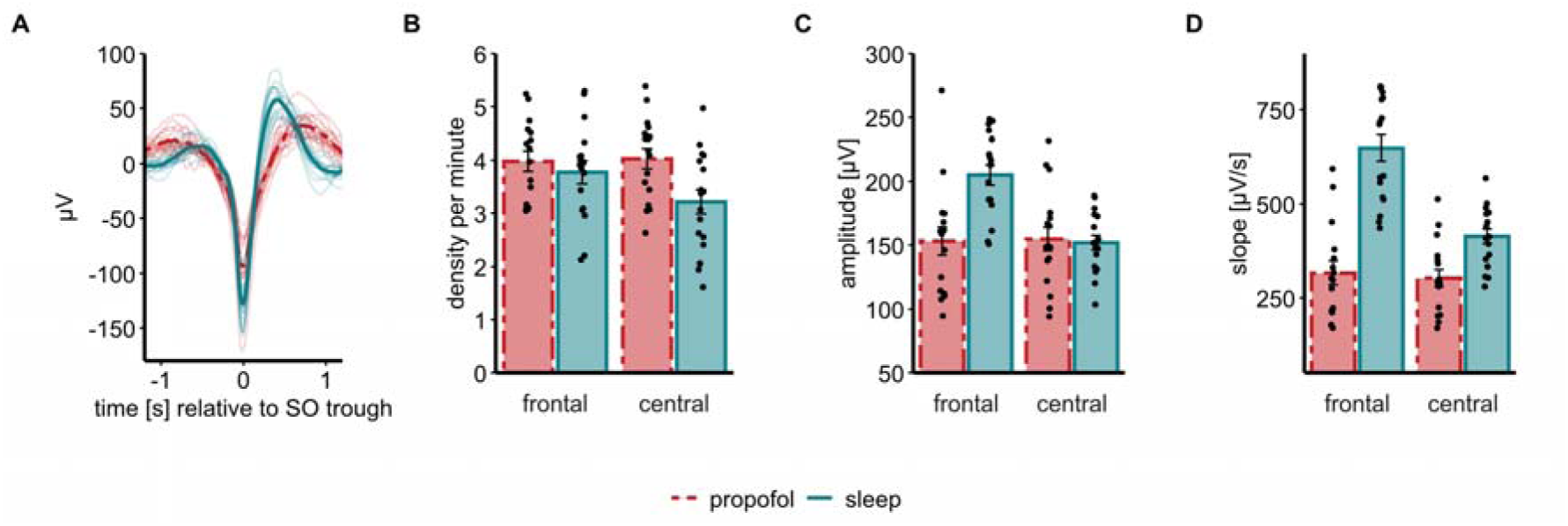
Slow oscillation characteristics during propofol-induced anesthesia and non-REM sleep. **A:** Grand-average frontal slow oscillation of the propofol (dark red, dashed lines) and sleep (teal, solid lines) group. Lighter traces depict individual participants. **B.** Slow oscillation density at frontal and central sites. **C.** Slow oscillation amplitude at frontal and central sites. **D.** Slow oscillation slope at frontal and central sites. B-D: depicted are individual data points, group means and standard errors.

Furthermore, we analyzed whether the groups differ in their SO trough-locked TFRs (Figure 6). Testing baseline-normalized TFRs against zero, revealed that both groups exhibited greater power during the SO trough in lower frequencies (sleep: cluster *p* < .001, propofol: cluster *p* = .003). However, only the sleep group showed an additional power increase in the fast spindle band during the SO upstate (cluster *p* = .001). Directly contrasting the baseline-normalized TFRs of the two groups revealed greater power changes in the sleep than the propofol group, especially in the fast spindle frequency range (cluster *p* = .001, Figure 6C). Complementing this picture, preferred phases of coupling were uniformly distributed in the propofol group (Rayleigh *z* = 0.66, *p* = .521) but non-uniformly distributed in the sleep group (Rayleigh *z* = 11.49, *p* < .001, Figure 6D).

**Figure 6.**
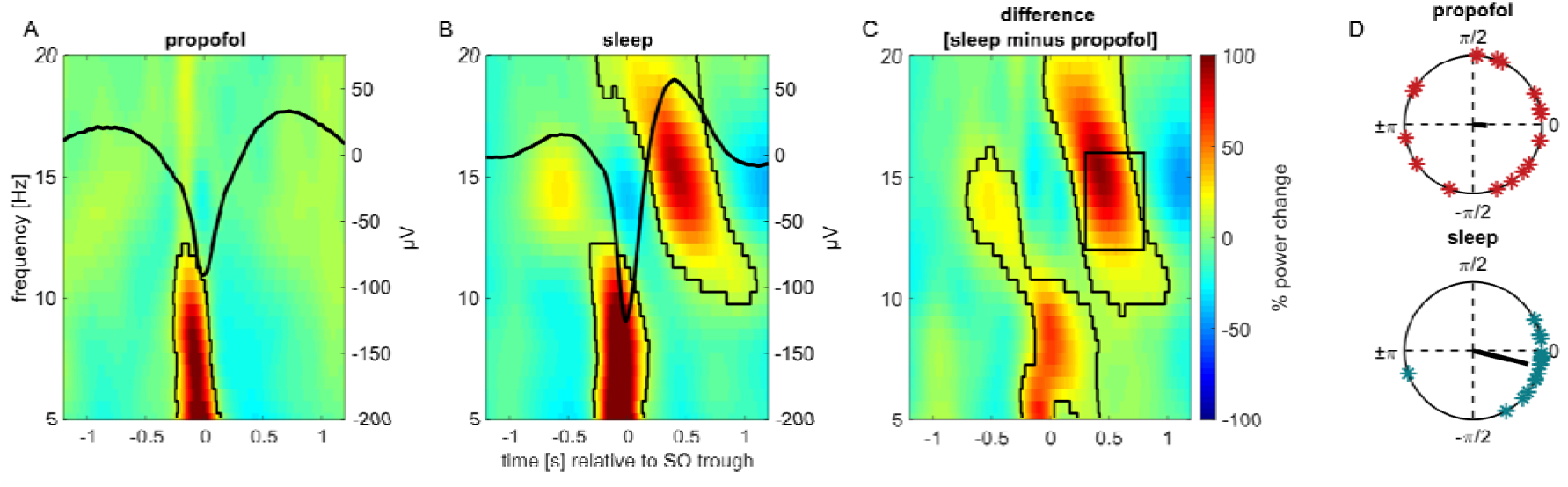
Frontal slow-oscillation trough-locked time-frequency representations and phase-amplitude coupling during propofol-induced anesthesia and sleep. **A:** Frontal time-frequency representation of SO events during propofol-induced anesthesia. **B.** Frontal time-frequency representation of SO events during non-REM sleep. A and B: Warm colors indicate greater power relative to baseline, and black contours indicate significant positive clusters. Average SO is superimposed **C.** Difference in baseline-normalized time-frequency representations between the propofol and sleep group. Warmer colors indicating greater changes in the sleep than the propofol group. Contours indicate significant positive clusters. **D.** Polar plots depicting the individual mean preferred phases of coupling during slow oscillations, separately for the propofol and sleep group.

Considering the role of non-REM sleep characteristics and their precise coupling in memory consolidation, we correlated SO characteristics and measures of SO-spindle coupling with changes in memory performance, i.e. post minus pre sleep/anesthesia. This did not reveal any significant associations, neither in the sleep nor in the propofol group (all *p* > .08).

## Discussion

In this study, we compared memory consolidation during natural sleep versus propofol-induced general anesthesia using the hippocampus-dependent virtual Morris water maze and a procedural memory task alongside concurrent EEG recordings. Our key findings reveal a behavioral dissociation: whereas sleep preserved or enhanced memory performance across the retention interval, propofol anesthesia led to significant deterioration in both tasks. Critically, while SOs occurred in both states, only sleep exhibited the canonical increase in spindle power during SO upstates, alongside the characteristic frontal-to-central decrease in SO amplitude. These results suggest that despite generating a superficially sleep-like oscillatory landscape, propofol nevertheless fails to support memory stabilization, likely due to a disrupted hippocampal-neocortical dialogue at systems level.

Based on the global GABAergic inhibition during anesthesia (Bao et al., 2023), its disruption of long-term potentiation in the hippocampus (Takamatsu et al., 2005; Wei et al., 2002) and the lack of sleep spindles coupled to SO upstates during anesthesia (Murphy et al., 2011), we expected propofol-induced general anesthesia to disrupt memory consolidation. Indeed, performance in the propofol group deteriorated across the retention interval comprising anesthesia independent of retention interval duration and age. This pattern contrasts sharply with sleep, where performance remained stable or improved, consistent with sleep’s established role in consolidating memories. Propofol has previously been shown to impair spatial memory consolidation in rats (Zhang et al., 2013) and hippocampal activity and memory in humans (Pryor et al., 2015).

Studies in humans suggest that the time between encoding and induction of anesthesia influences the level of memory impairment (Iggena et al., 2022; Moon et al., 2020). Using a variation of the virtual water maze task, Iggena et al. (2022) found worse spatial memory performance when anesthesia was induced directly after encoding compared to induction 60 minutes after encoding or compared to a wake control group. Although our surgical setting introduced variable delays between encoding and anesthesia (as well as post-anesthesia recovery), retention interval duration did not explain the group difference, suggesting a robust amnestic effect even under ecologically valid conditions. While our study showed diminished memory performance after anesthesia in a spatial and procedural memory task, a recent study found that gist-based memory might be less affected by propofol (Risse et al., 2025), highlighting that non-hippocampal gist-like processing may resist propofol’s effects, and further underscoring our tasks’ sensitivity to hippocampus-dependent consolidation. Together, this supports the idea that both procedural and declarative memory consolidation during sleep relies on the hippocampus.

During sleep, the repeated reactivation of newly encoded information in hippocampal and neocortical sites in concert with the precise coupling of SOs, spindles and ripples promotes memory consolidation (Brodt et al., 2023). The precise coupling of SOs and fast spindles has repeatedly been associated with sleep dependent memory consolidation in different tasks (Ng et al., 2025). This was similarly demonstrated for the water maze task (Bastian et al., 2022) and the mirror tracing task (Mikutta et al., 2019), the tasks that were used in the current study. Our polysomnographic approach allowed direct quantification of these parallels and their critical divergences. First, we found that the oscillatory peak in the power spectra during non-REM sleep reflected the occurrence of sleep spindles. Frontal sleep spindles are usually slower than centro-parietal spindles (Cox et al., 2017), which was similarly seen here with a smaller sigma peak at frontal than central sites. Contrarily, propofol-induced anesthesia was characterized by continuous alpha activity (Purdon et al., 2013). Secondly, both states were characterized by the occurrence of SOs. For each of the assessed SO measures, we saw a frontal to central decrease in the sleep group while SO density, amplitude and slope were similar at both sites in the propofol group. Our results in the sleep group reflect the frontal predominance of SOs and the anterior to posterior decrease in amplitude and slope during non-REM sleep (Riedner et al., 2007). However, contrary to Murphy et al. (2011) who did not find any topographical differences between sleep and propofol in any of the SO measures, we found higher SO density centrally and lower amplitude frontally as well as lower slopes at both sites in the propofol compared to the sleep group suggesting less synchronized, possibly fragmented cortical involvement. Most importantly, and in line with Murphy et al. (2011), only SOs during non-REM sleep but not propofol were associated with an increase in spindle activity during the SO upstate in our study. Together with the superior effect of sleep on spatial and procedural memory consolidation our results suggest the lack of SO-spindle coupling during anesthesia as a possible reason for impaired consolidation.

However, we did not find any association between memory consolidation and SO-spindle coupling during sleep. Contrary to Mikutta et al. (2019) where participants performed each three encoding and three retrieval trials in the mirror tracing task, our participants performed one trial each. Thus, although most of the participants in the sleep group showed an improvement in their performance and consolidation was better than in the propofol group, it is conceivable that reaching a certain pre-sleep performance level might have particularly benefited from the precise nesting of spindles in the SO upstate (Cross et al., 2025). However, we likewise did not find an association between changes in memory performance and SO-spindle coupling in the spatial task, although there, participants performed nine learning trials, thus improving their performance level pre offline consolidation. This absence of an association in the sleep group might also be owed to a lack of power due to the small sample size.

Propofol’s amnesic profile may arise from multi-level disruptions. It inhibits hippocampal long-term potentiation in CA1 via enhanced GABA-A-phasic and tonic currents, directly impairing synaptic plasticity critical for consolidation (Takamatsu et al., 2005; Wei et al., 2002). Computational models suggest that baseline waxing and waning thalamic spindling is replaced by a sustained alpha rhythm during propofol induction (Soplata et al., 2017). Furthermore, information integration is impaired through propofol-induced disruption of thalamocortical connectivity (Liu et al., 2013). These mechanisms converge on mnemonic information processing: while during sleep the triple coupling of SOs, spindles and ripples orchestrates hippocampo-neocortical transfer, propofol disrupts this “dialogue” at multiple nodes - hippocampal plasticity, thalamic pacemaking, and cortical integration. Together our results indicate impaired memory consolidation during propofol-induced general anesthesia compared to natural sleep. A plausible framework for the diminished memory consolidation under propofol-induced anesthesia is the disruption of hippocampal-neocortical dialogue relayed by SO-spindle coupling during non-REM sleep.

## Author contributions

T.B., A.H., and A.F. conceptual development and design of the study. M.D., F.H. A.F. J.R. and S.P. collected the data E.-M. K., A.H.: formal analysis, data interpretation, writing – original draft F.D.W., R.S. contributed to data analysis and interpretation J.B., S.B., T.B. supervision, writing – review & editing All authors contributed to the final version of the manuscript and approved its submission.

## Data and code availability

The data that support the findings of this study are available on request from the corresponding author. The data are not publicly available due to privacy or ethical restrictions.

## Acknowledgements

We thank all participants for taking part in this study. This work was supported by the German Research Foundation (DFG) FOR 5434.

## Declaration of interests

The authors declare no competing interests.

## Notes

### Competing Interest Statement

The authors have declared no competing interest.

