## Supporting Information for "Natural sleep, but not propofol-induced anesthesia preserves spatial and procedural memory consolidation"

‡ these authors contributed equally

*Corresponding author

### Supplementary methods

#### Propofol-induced general anesthesia

*Induction and maintenance of anesthesia*

Anesthesia was performed according to the local standard operating procedures and without study-specific amendments. Standard monitoring including pulse oximetry, non-invasive blood pressure measurement and ECG were established before anesthesia induction. After preoxygenation, general anesthesia was induced via target-controlled infusion (TIVA/TCI; B. Braun Melsungen AG, Melsungen, Germany) and infusion of remifentanil (0.2–0.7 µg/kg/h; GlaxoSmithKline, Munich, Germany). The effective biological half-life of remifentanil is 3 to 10 min. Plasma and effect-site concentrations of propofol were estimated by means of a 3-compartment pharmacokinetic model. The mean calculated cerebral target concentration was 2.53 ± 0.11 µg/ml. After loss of consciousness, endotracheal intubation was facilitated by intravenous rocuronium (0.6 mg/kg; Essex Pharma, Munich, Germany). Ventilation was pressure controlled using an S/5TM ADU anesthesia device (GE Healthcare Finland Oy, Helsinki, Finland). Depth of sedation was controlled regularly from induction to recovery by means of the Ramsay Sedation Scale (Ramsay et al., 1974). During surgery, the depth of anesthesia was controlled by adjusting the infusion rate of remifentanil and the target concentration of propofol as needed. Blood pressure, heart rate, remifentanil infusion rate, propofol plasma target concentration (Cpt) and calculated cerebral propofol concentration (Ce) and BIS were documented in 5 min intervals throughout the procedure. In addition, special events such as patient movements or the administration of additional medication were noted.

*Recovery from Anesthesia*

The administration of remifentanil and propofol was discontinued shortly before or after the end of the operation after assuring neuromuscular recovery by neuromuscular monitoring. For postoperative pain therapy, the patients received 1 g paracetamol (Rotexmedica GmbH, Trittau, Germany) intravenously before the anesthetics were withdrawn. If necessary, the patients received further analgesics. After the patients regained consciousness and protective reflexes returned, extubation was performed. Postoperative monitoring of the patients was ensured in the recovery room. Prior to delayed recall, intensity of pain was assessed using a numerical rating scale (0=no pain, 10 = worst imaginable pain, 3.92 ± 0.63).

#### Collection of cortisol levels

As stress has been found to impair memory performance (De Quervain et al., 2000; Wagner et al., 2005), cortisol in saliva was assessed by collecting samples on the morning of the operation and immediately stored at -25°C until analysis via immunoassay (Immulite, Siemens Healthcare, Erlangen, Germany, intra-assay coefficient of variation 6.8%, inter-assay coefficient of variation 9.9%). Average preoperative cortisol levels were 1.26 ng/ml ± 0.35 (range 0.03 – 4.7 ng/ml, normal circadian matched range 0.04 – 4.67 ng/ml) showing physiological morning cortisol levels.

**Electroencephalographic recording and analyses**

EEG recordings were sampled at 128 Hz in the propofol group and 256 Hz in the sleep group, respectively. In both groups, Cz was the recording reference. Offline, the EEG was re-referenced to the average potential of the left and right mastoid and down sampled to 128 Hz. In both groups, EEG was recorded from F3, F4, C3, and C4. In the sleep group, O1, and O2 were additionally recorded. In addition, electrooculogram (EOG) and electromyogram (EMG, sleep group only) were assessed.

EEG and EOG signals were filtered between 0.3 and 35.0 Hz and EMG between 2 Hz and 128.0 Hz. In the sleep group, sleep stages were determined in 30-second epochs based on standard criteria of the American Academy of Sleep Medicine (Berry et al., 2017). Total sleep time (TST), time spent in stage 1 (N1), stage 2 (N2), slow wave sleep (SWS/N3), and REM sleep as well as wake after sleep onset (WASO) were determined. In the propofol group, loss (recovery) of consciousness was determined based on the occurrence (vanishing) of alpha and delta activity (Purdon et al., 2013). Artefactual epochs were marked during scoring and excluded from further analyses.

*Power Spectral activity*

Power spectra of non-REM sleep and propofol-induced anesthesia were calculated using the FieldTrip toolbox (Oostenveld et al., 2011) in MATLAB 2023b. The artefact-free data was epoched into 10 second segments and power calculated using fast fourier transformation with discrete prolate spheroidal sequences tapers (9 tapers) and 0.5 Hz smoothing (*mtmfft*). For each participant, the resulting spectra were averaged across frontal (F3/F4) and central (C3/Cz/C4) channels and then subjected to the python-based fooof (fitting oscillations & one over f) toolbox (Donoghue et al., 2020) in order to obtain aperiodic (1/*f* like characteristics) and periodic components in a frequency range between 0.5 and 30 Hz using the fixed mode. Further settings included a peak width limit between 0.5 and 12 Hz, a maximum number of three detected peaks, a minimum peak height of 0 and a peak threshold of 2 SDs. For each participant the dominant frequency between 0.5 and 30 Hz, i.e. frequency peaks in the "aperiodic corrected" power spectra, was obtained.

*Slow Oscillation detection and cross-frequency coupling*

Detection of SOs during non-REM sleep and anesthesia were done using the MATLAB-based toolbox SleepTrip (SCR_017318), with algorithms based on Mölle et al. (2002). After applying a 0.3 Hz high- and a 4 Hz low-pass filter, all consecutive positive-to-negative zero crossings were marked as potential SOs in the signal. Only SOs with a frequency between 0.5 and 1.25 Hz, an amplitude 1.25 times greater and a trough peak potential 1.25 times lower than the average of the potential SOs within a channel of a participant were considered as SOs. For each channel and participant, the SO density (# per minute), the amplitude (µV, maximum trough to peak potential), the ascending slope (µV/s, ratio between the trough potential and the time delay to the following up-zero crossing; Kurth et al., 2010) and duration (time between the two positive-to-negative zero crossings) were obtained and averaged across frontal and central channels. Since the EEG recordings during surgery were quite noisy, the detected events were visually verified in the continuous EEG for each participant.

Further analyses focused on the frontal channels (F3/F4). For each channel, the ±3 seconds around each SO trough were extracted. Time-frequency representations (TFR) were calculated for the SO trough-locked data in a frequency range of 5 to 20 Hz in 0.5 Hz steps using the FieldTrip toolbox. Morlet wavelets with a width linearly increasing from 4 to 12 were used. TFRs were normalized by calculating the relative power change with respect to the mean power within each frequency bin in the ±1.5 s window around the SO trough.

Additionally, we calculated the Synchronization Index (SI, Cohen, 2008) for each extracted SO time series, to determine the amount of power modulations in the spindle frequency band (12-16 Hz) by SO activity (0.5-1.25 Hz). See Kurz et al. (2021) and Staresina et al. (2015) for further details. For each participant and channel, we calculated the mean preferred phase of coupling and the vector length as a measure of coupling consistency using the CircStat toolbox in MATLAB (Berens, 2009). Within each participant and across groups, we used the Rayleigh Test to determine (non-)uniformity of the preferred phases of coupling.

We refrained from the detection of spindle events because the continuous alpha activity during propofol-induced anesthesia could lead to ambiguous or false-positive spindle detections in the propofol group. Thus, we opted for the cross-frequency coupling described above. Although the spindle frequency partially overlaps with propofol-related alpha activity, the analysis does not assume the presence of discrete spindle events. Rather, it quantifies whether SOs systematically modulate 12–16 Hz frequencies, as expected for sleep-related SO-spindle coupling but not for continuous propofol-induced alpha activity.

**Table S1.** Neuropsychological Assessment (M ± SEM)

| | Propofol *n* = 16 | Sleep *n* = 17 | *z*/*t* | *p* |
|---|---|---|---|---|
| RAVLT sum | 54.50 ± 2.32‡ | 59.70 ± 1.77 | 1.76 | .089 |
| RAVLT retention | 13.38 ± 0.68 | 13.41 ± 0.44 | -0.76 | .460 |
| RAVLT delayed | 13.50 ± 0.62 | 13.35 ± 0.47 | -0.59 | .568 |
| digit span forward | 8.88 ± 0.58 | 7.94 ± 0.61 | -1.11 | .274 |
| digit span backward | 6.88 ± 0.53 | 8.18 ± 0.68 | 1.51 | .141 |
| word fluency lexical† | 8.50 ± 0.87 | 9.27 ± 0.45 | 0.78 | .448 |
| word fluency categories‡ | 10.68 ± 1.08 | 14.38 ± 0.51 | 3.11 | .007 |
| TMT A | 24.56 ± 2.24 | 21.00 ± 1.35 | -1.48 | .143 |
| TMT B | 62.31 ± 5.81 | 54.12 ± 4.58 | -1.28 | .207 |

RAVLT: Rey-Auditory Verbal Learning Test; TMT = Trail Making Test; † n propofol = 10; ‡ n propofol = 11

**Table S2.** Virtual Water Maze Task (M ± SEM)

| | latency | | path length/pool diameter | | relative dwell time | |
|---|---|---|---|---|---|---|
| | Propofol *n* = 13 | Sleep *n* = 17 | Propofol *n* = 13 | Sleep *n* = 17 | Propofol *n* = 13 | Sleep *n* = 17 |
| LT 1 | 60.08 ± 7.40 | 40.43 ± 1.92 | 1.37 ± 0.09 | 1.21 ± 0.03 | 0.79 ± 0.06 | 0.75 ± 0.07 |
| LT 2 | 34.72 ± 5.43 | 68.93 ± 18.24 | 1.06 ± 0.19 | 2.40 ± 0.73 | 0.88 ± 0.03 | 0.72 ± 0.07 |
| LT 3 | 19.37 ± 4.33 | 35.62 ± 14.15 | 0.41 ± 0.10 | 1.23 ± 0.51 | 0.99 ± 0.01 | 0.92 ± 0.04 |
| LT 4 | 46.36 ± 7.77 | 48.12 ± 7.79 | 1.31 ± 0.16 | 1.86 ± 0.30 | 0.94 ± 0.01 | 0.76 ± 0.05 |
| LT 5 | 34.08 ± 6.16 | 34.32 ± 5.06 | 1.28 ± 0.23 | 1.60 ± 0.24 | 0.90 ± 0.02 | 0.81 ± 0.03 |
| LT 6 | 13.92 ± 2.88 | 36.69 ± 7.41 | 0.32 ± 0.05 | 1.46 ± 0.37 | 1.00 ± 0.00 | 0.86 ± 0.05 |
| LT 7 | 20.40 ± 2.79 | 40.00 ± 13.83 | 0.73 ± 0.06 | 1.68 ± 0.52 | 0.93 ± 0.01 | 0.83 ± 0.05 |
| LT 8 | 24.16 ± 3.15 | 45.47 ± 8.46 | 0.87 ± 0.07 | 2.00 ± 0.49 | 0.94 ± 0.004 | 0.81 ± 0.06 |
| LT 9 | 13.55 ± 2.22 | 29.76 ± 7.69 | 0.33 ± 0.03 | 1.22 ± 0.42 | 1.00 ± 0.00 | 0.89 ± 0.05 |
| RT 1 | 53.10 ± 9.17 | 41.26 ± 5.33 | 1.64 ± 0.21 | 1.63 ± 0.21 | 0.72 ± 0.06 | 0.83 ± 0.03 |

LT: learning trial; RT: retrieval trial

**Table S3.** Mirror Tracing Task (M ± SEM)

| | Propofol<br>$n$ = 16 | Sleep<br>$n$ = 17 |
|---|---|---|
| errors | | |
| pre | 26 ± 4.35 | 33 ± 6.53 |
| post | 21 ± 5.94 | 14 ± 3.03 |
| Duration (sec)† | | |
| pre | 112 ± 16 | 110 ± 9.71 |
| post | 75 ± 6.39 | 69 ± 4.94 |

† n propofol = 14
